# Lack of apolipoprotein A1 impairs optimal regulatory T cell homeostasis at steady state due to impaired IL-2 signaling

**DOI:** 10.1101/751107

**Authors:** Dalia E. Gaddis, Runpei Wu, John S. Parks, Mary G. Sorci-Thomas, Catherine C. Hedrick

**Author notes:** Corresponding Author: Catherine C. Hedrick, Ph.D., La Jolla Institute for Immunology, 9420 Athena Circle, La Jolla, CA 92037.

## Abstract

Apolipoprotein A1 (ApoA1), the major constituent of the high-density lipoprotein (HDL) molecule, exhibits anti-inflammatory properties. Our laboratory has previously shown that ApoA1 protects against switching of regulatory T (Treg) cells to atherogenic T follicular helper cells in Western diet-fed mice. However, the role of ApoA1 in modulating Treg cell homeostasis in the absence of atherosclerosis remains uncharacterized. Here, we show that ApoA1 is required for normal Treg cell homeostasis and functioning at steady state. Specifically, lack of ApoA1 decreased the numbers of both natural and induced Treg cells and also lowered Treg cell-based homeostatic proliferation and suppressive functions. Importantly, these changes occurred without affecting other T cell populations. Finally, we determined that the observed phenotypes were caused by changes to cholesterol content and reduced interleukin-2 (IL-2) receptor signaling in ApoA1-deficient Treg cells. Overall, our results show that ApoA1-HDL is necessary for Treg cell homeostasis and functioning.

## INTRODUCTION

Elevated, sustained levels of high-density lipoprotein (HDL) are associated with an increased risk for atherosclerosis and cardiovascular disease. Apolipoprotein AI (ApoA1) is the major protein component of HDL, accounting for approximately 70% of the molecule. ApoA1 is produced in the liver and is essential for the normal function of HDL (1). Through ATP-binding cassette transporter A1 (ABCA1), ApoA1 promotes cholesterol efflux to form HDL (2–4). ApoA1 thus aids in the excretion of cholesterol, a process that is beneficial for overall cardiovascular health.

Both HDL and ApoA1 have been shown to have anti-inflammatory properties (5). For example, ectopic expression of ApoA1 in macrophages reduces immune infiltration in the aorta and protects against atherosclerosis in a cardiovascular disease mouse model (6). Additionally, administration of ApoA1 reduces skin inflammation, T cell proliferation, and the ratio of effector T cells to regulatory T cells in mice with loss of both ApoA1 and the low-density lipoprotein receptor (LDLR) (7, 8). More recently, our laboratory has found that ApoA1 prevents the switch of protective, regulatory T cells to atherogenic, inflammatory T follicular helper cells, and also reduces B cell germinal center formation and activation, in an atherogenic mouse model (9). Importantly, these findings extend the anti-inflammatory effects of ApoA1 beyond cardiovascular disease. Indeed, ApoA1 attenuates neuroinflammation in a mouse model of Alzheimer’s disease (10) and reduces lymphocyte activation and autoimmunity in a lupus mouse model (11). Similarly, a model of rheumatoid arthritis shows reduction of autoimmune responses with the presence of ApoA1 (12).

ApoA1 decreases the differentiation and maturation of human dendritic cells and thus dampens T cell activation (13), suggesting that ApoA1 can influence T cell responses at steady state. However, the function of ApoA1 in the immune system during steady state has yet to be fully determined. Thus, we sought to determine the involvement of ApoA1 on T cell immune homeostasis during steady state and the consequences of ApoA1 deficiency on T cell development.

## MATERIALS AND METHODS

### Mice

Breeding pairs of C57BL/6 (B6), *Apoa1*^−/−^, and CD45.1 mice were purchased from The Jackson Laboratory (Bar Harbor, Maine; Stock numbers 000664, 002055, and 002014, respectively). For mice with ABCA1 loss specifically in T cells, bone marrow chimeras were generated from bone marrow cells collected from *Abca1*^flox/flox^ and *Abca1*^flox/flox^;*Lck*-Cre^+/−^ mice (both from John S. Parks, Wake Forest University). Recipient B6 mice were irradiated with 550 rads twice, four hours apart. Approximately 1×10^7^ bone marrow cells were retro-orbitally injected into recipient mice. Recipient mice were then allowed to reconstitute for 10 weeks before they were sacrificed and their peri-aortic lymph nodes (PaLNs) were analyzed for Treg cells. Female mice were used for experiments at 6-10 weeks of age. All mice were fed a standard chow diet containing 0 % cholesterol and 5 % calories from fat (Pico lab, #5053, Saint Louis, MO). Mice were housed in a pathogen-free animal facility of La Jolla Institute for Immunology (LJI). All experiments followed the guidelines of the LJI Animal Care and Use Committee and approval for use of rodents was obtained from LJI according to criteria outlined in the Guide for the Care and Use of Laboratory Animals from the National Institutes of Health.

### Flow cytometry

PaLNs, other peripheral lymph nodes, thymii, or spleens were passed through 40 μm cell strainers after homogenization. For spleens and thymii, RBCs were lysed with 1X RBCs lysis buffer (Biolegend, San Diego, CA). All staining was done on ice for 30 minutes unless otherwise indicated. Antibodies were purchased from eBioscience, Biolegend or BD Bioscience (San Diego, CA) unless otherwise indicated. Samples were stained with Live Dead fixable dye (Thermo Fisher, Carlsbad, CA). Cells were surface stained with antibodies against CD4 (clone RM4-4), TCRβ (clone H57-597), CD25 (clone PC61), membrane-bound latent TGFβ (clone TW7-16B4), Nrp1 (clone N43-7; MBL International, Woburn, MA), and CD45.1 (clone A20; for suppression assay) in cold FACS buffer (2 % BSA, 0.01 % sodium azide in PBS). Cells were fixed and permeabilized with Foxp3 Staining Buffer Set (eBioscience), then stained with antibody against Foxp3 (clone FJK-16S) and/or Caspase 3 (BD Biosciences) in permeabilization buffer. Filipin III (Cayman Chemical, Ann Arbor, MI) was used to stain for cholesterol content, as recommended by the manufacturer’s instructions. For pSTAT5, cells were stimulated with recombinant human IL-2 (100 U/ml) (Peprotech, Rocky Hill, NJ) for 45 min at 37°C. Cells were then surface stained as previously described. Cells were washed and fixed with 1X Lyse/Fix buffer (BD Biosciences) for 10 min at 37°C. Cells were then washed and chilled on ice and permeabilized with pre-chilled BD phosphoflow Perm Buffer III for 30 min (BD Biosciences), followed by staining for Foxp3 and rabbit pSTAT5 (Cell Signaling Technologies, Beverly, MA), and then by secondary anti-rabbit-AF647 (eBioscience). Samples were acquired using an LSRII flow cytometer (BD, Bioscience, San Diego, CA) and data were analyzed using Flowjo Ver 9.8 and 10.0.08 (Tree Star, Ashland, OR).

### *In vivo* proliferation assay

B6 and *Apoa1*^−/−^ mice were given intraperitoneal injections of 1 mg of BrdU. PaLNs were then harvested 36 hours after injection. PaLN cell suspensions were stained as explained above. Cells were then DNase treated and stained for BrdU as recommended by the BrdU Flow Kit (BD Biosciences).

### Suppression assay

Treg cells (CD4^+^CD25^+^) were sorted from spleens and peripheral LNs of B6 and *Apoa1*^−/−^ mice. Naïve effector (CD4^+^CD25^−^CD44^lo^CD62L^hi^) cells were sorted, using FACSAria (BD Biosciences), from spleens and peripheral LNs of CD45.1 mice and were labeled with cell trace violet (CTV) (Thermo Fisher) according to the manufacturer’s instruction. Feeder cells were prepared from spleens of B6 mice, with CD4 T cells depleted using CD4^+^ magnetic beads (Miltenyi Biotec, Auburn, CA), and then irradiated with 3000 rads. Feeders were cultured with naïve effector T cells at a ratio of 3:1 in U-shaped 96 well plates. Sorted Treg cells from B6 or *Apoa1*^−/−^ mice were added at a ratio of 1:2, 1:4, or 1:8. Cells were then stimulated with soluble αCD3 antibody at 2.5 μg/ml. All cells were cultured in RPMI media supplemented with 10 % FCS, L-glutamine, penicillin, streptomycin, and β-mercaptoethanol. Cells were harvested 4 days following stimulation and the percentage of effector CD45.1 cells that diluted CTV was determined.

### *In vitro* Treg induction

Naïve CD4 T cells were sorted as described above or using EasySep™ Naïve CD4 T cell Isolation Kit (STEMCELL Technologies, Inc., Vancouver, Canada). Equal numbers of cells in serum-free, CST OpTmizer™ T cell expansion media (Thermo Fisher), supplemented with L-glutamine, penicillin, streptomycin, and β-mercaptoethanol, were incubated in αCD3-coated (1 μg/ml) plates. Soluble αCD28 (1 μg/ml) and TGFβ (2 ng/ml) were added to the wells. Cells were incubated for 2 days (for IL-2 detection) or 4 days (for induced Treg staining).

### ELISA

Supernatants from *in vitro* Treg induction were collected following 2 days of stimulation and frozen at −80°C until analyzed. IL-2 ELISAs were performed using Ready-Set-Go ELISA kits (eBioscience) according to manufacturer’s instruction.

### Statistical analyses

All results are expressed as means (± SEM). Results were analyzed by unpaired, two-tailed Student’s *t*-tests. *p* values of *<* 0.05 were considered to be statistically significant. Statistical analyses were performed using GraphPad Prism software version 6 (GraphPad Software, Inc.).

## RESULTS

### ApoA1 is required for optimal Treg cell homeostasis in steady state

To determine the role of ApoA1 in Treg cell development during steady state, we compared Treg cells in the peri-aortic lymph nodes (PaLNs) of *Apoa1*^−/−^ mice to wild-type, C57BL/6 (B6) mice. We found a decrease in the percentages and numbers of CD4^+^CD25^+^Foxp3^+^ Tregs (~ 27 % and 50 %, respectively) in *Apoa1*^−/−^ mice compared to B6 controls (**Figure 1A**). These changes were accompanied by an approximately 50 % decrease in the expression of latent, membrane-bound transforming growth factor β (TGFβ) on Treg cells (**Figure 1B**). There was also an almost 50 % decrease in the numbers of naturally occurring Treg cells in *Apoa1*^−/−^ mice, as indicated by a decrease in the number of Treg cells positive for neuropilin1 (NRP1), a marker for thymus-derived Treg cells (14, 15) (**Figure 1C**). Furthermore, we found no changes in effector or memory CD4 T cell development (**Figure 1D**) or the ability of CD4 T cells to produce interferon (IFN)-γ or interleukin (IL)-17 (**Figure 1E**) in *Apoa1*^−/−^ mice. Finally, the observed reduction in peripheral Treg cells was not due to a decrease in the percentages or numbers of Treg cells in the thymus (**Figure 1F**), suggesting a defect specifically in Treg cell homeostasis.

**Figure 1.**
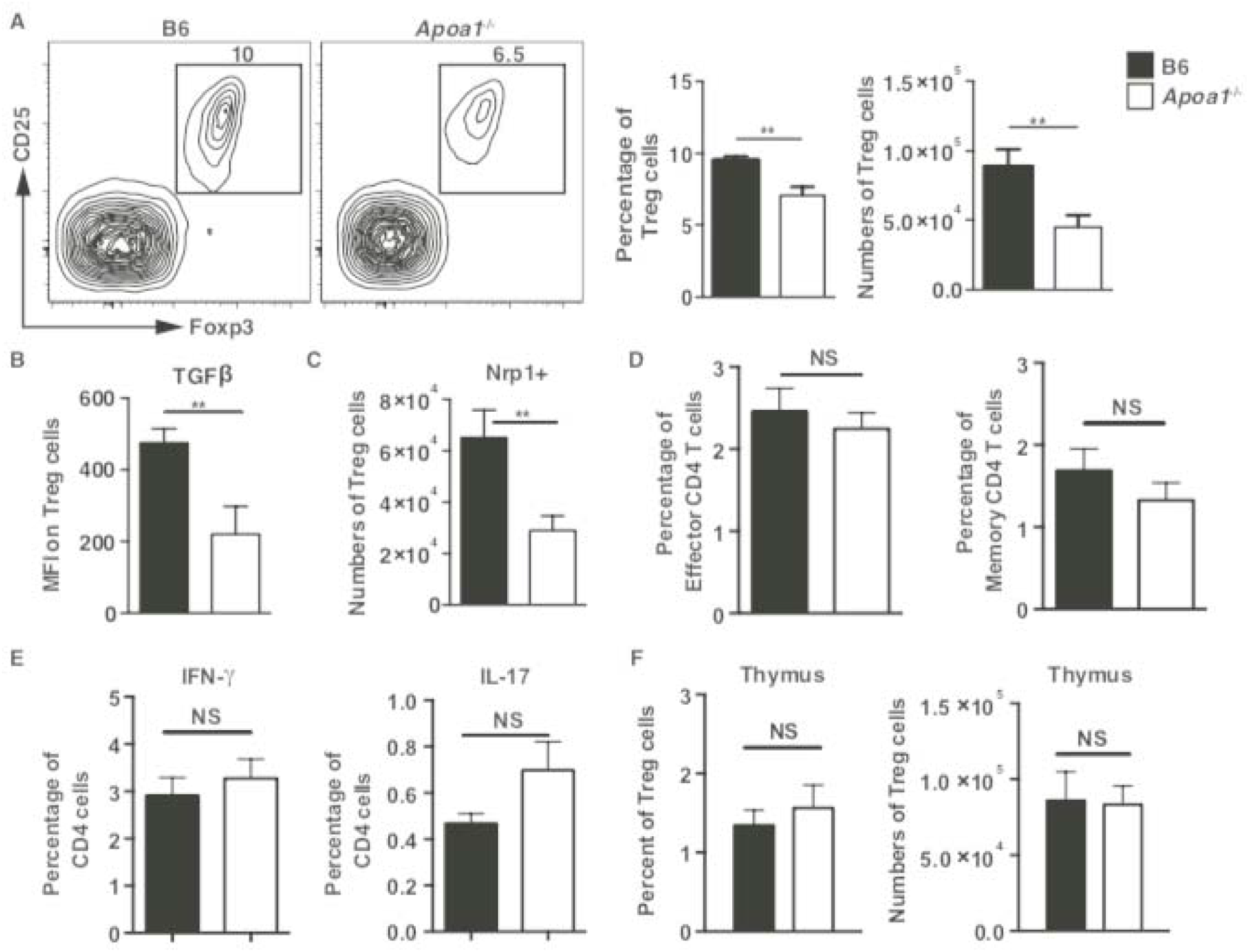
In steady state, ApoA1 is required for optimal Treg cell development. **(A-C)** Peri-aortic LNs (PaLNs) from B6 and *Apoa1*^−/−^ mice were examined for **(A)** the percentages and numbers of Treg (CD4^+^CD25^+^Foxp3^+^) cells, **(B)** the expression of membrane-bound latent TGFβ (as geometric mean fluorescence intensity), **(C)** the numbers of Nrp1^+^ Tregs by flow cytometry, and (**D**) the percentages of effector (CD44hiCD62Llo) and memory (CD44hiCD62Lhi) CD4 T cells. (**E**) Cells from the above mice were also examined for expression levels of IFN-γ and IL-17, following PNA/Ionomycin stimulation for 5 hours, via flow cytometry. (**F**) Treg cell percentages and numbers in the thymus of B6 and *Apoa1*^−/−^ mice. Results are expressed as means (± SEM), for three independent experiments (n = 13-15), \*\**p* < 0.01, NS = no significant differences.

### ApoA1-deficient Treg cells show reduced proliferative and suppressive capacities

We next sought to examine if the decreased numbers of Treg cells in *Apoa1*^−/−^ mice were due to a defect in their homeostatic, proliferative capacity. Thus, BrdU injection in B6 and *Apoa1*^−/−^ mice revealed an approximately 30% decrease in the proliferative capacity of *Apoa1*^−/−^ Treg cells in PaLNs, compared to B6 cells (**Figure 2A**). Moreover, this defect was only detected in Treg cells and not in naïve or memory CD4 T cells (**Figure 2A**). Treg cells in PaLNs of *Apoa1*^−/−^ mice also showed no differences in their rates of apoptosis, as measured by the frequency and number of caspase3^+^ Treg cells (**Figure 2B**). These data suggest that the reduction of the number of Treg cells in *Apoa1*^−/−^ mice is not due to an increase in Treg cell apoptosis but rather a defect in their proliferative capacity.

**Figure 2.**
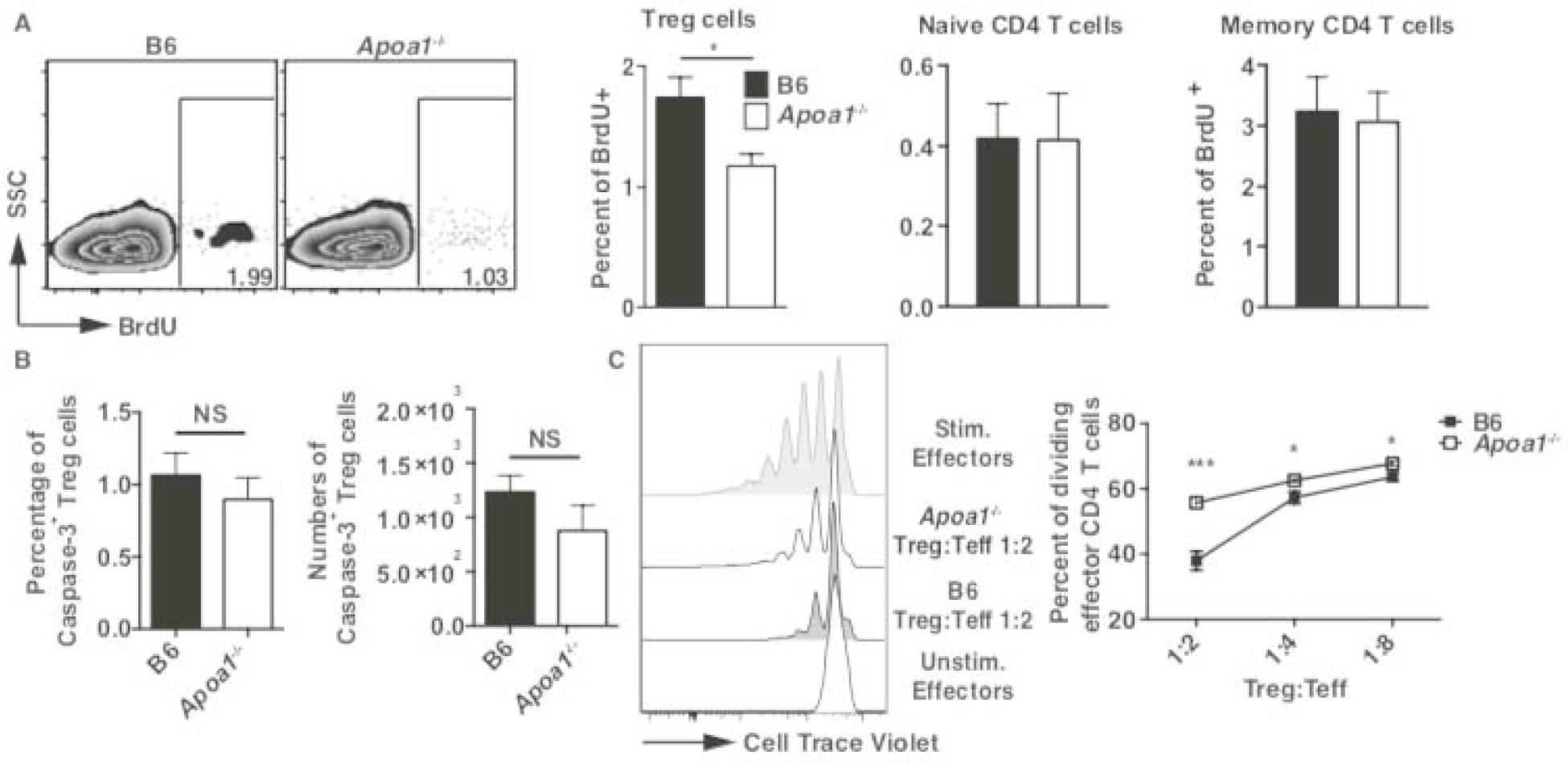
ApoA1 is required for optimal regulatory T cell proliferation and function. **(A)** B6 and *Apoa1*^−/−^ mice were injected i.p. with 1 mg of BrdU, PALNs were harvested 36 hours later and cells were stained for BrdU incorporation in T cells. Percentages show BrdU^+^ cells in Treg, naïve, and memory CD4 T cells. **(B)** Caspase-3^+^ staining of Treg cells in PaLNs from B6 and *Apoa1*^−/−^ mice, showing both percentages and numbers of Caspase-3^+^ cells. **(C)** Sorted Treg cells (CD4^+^CD25^+^) from spleens and peripheral LNs of B6 and *Apoa1*^−/−^ were cultured with CD4-depleted feeder cells from spleens of B6 mice and increasing numbers of naïve effector CD45.1 CD4 T cells (CD25^−^CD44^lo^CD62L^hi^) that were labeled with Cell Trace Violet (CTV). Cells were stimulated with soluble αCD3 for 4 days, harvested, and effector cells (CD4^+^CD45.1^+^) were examined for the dilution of CTV by flow cytometry. Results are expressed as the mean (± SEM) from two independent experiments (n = 9) **(A)**, the mean (± SEM) from two independent experiments (n = 8) **(B)**, and the mean (± SEM) of triplicate wells from one of three independent experiments **(C)**, \**p* < 0.05 and \*\**p* < 0.01.

To determine whether ApoA1 had any direct effect on Treg cell function, we tested the ability of Treg cells from *Apoa1*^−/−^ mice to suppress the proliferation of effector CD4 T cells. We observed that *ApoA1*^−/−^ Treg cells had a lower suppressive capacity than B6 Treg cells, (~ 55% versus ~ 38% of effector cells dividing in a 2:1 Teff:Treg culture, respectively) (**Figure 2C**). These results suggest that ApoA1 is critical for homeostatic proliferation and functioning of peripheral Treg cells during steady state.

### Loss of ApoA1 increases Treg cell cholesterol content

Since ApoA1 facilitates cholesterol efflux, we hypothesized that Treg cells from *Apoa1*^−/−^ mice would display altered cholesterol processing. Thus, we measured levels of intracellular cholesterol by filipin III staining and observed a significant increase in the fluorescence intensity of filipin III in *Apoa1*^−/−^ Treg cells compared to those from B6 mice (**Figure 3A**). ApoA1 couples the cholesterol transporter ATP-binding cassette transporter A1 (ABCA1) to allow efflux of cellular cholesterol. We therefore tested if mice with T cell-specific deletion of ABCA1 displayed defects in Treg development. We found a ~ 20 % decrease in the percentages of Treg cells in mice lacking ABCA1 in T cells (**Figure 3B**). Together, these data suggest that defects in cholesterol transport by either loss of ApoA1 or ABCA1 impairs Treg cell homeostasis.

**Figure 3.**
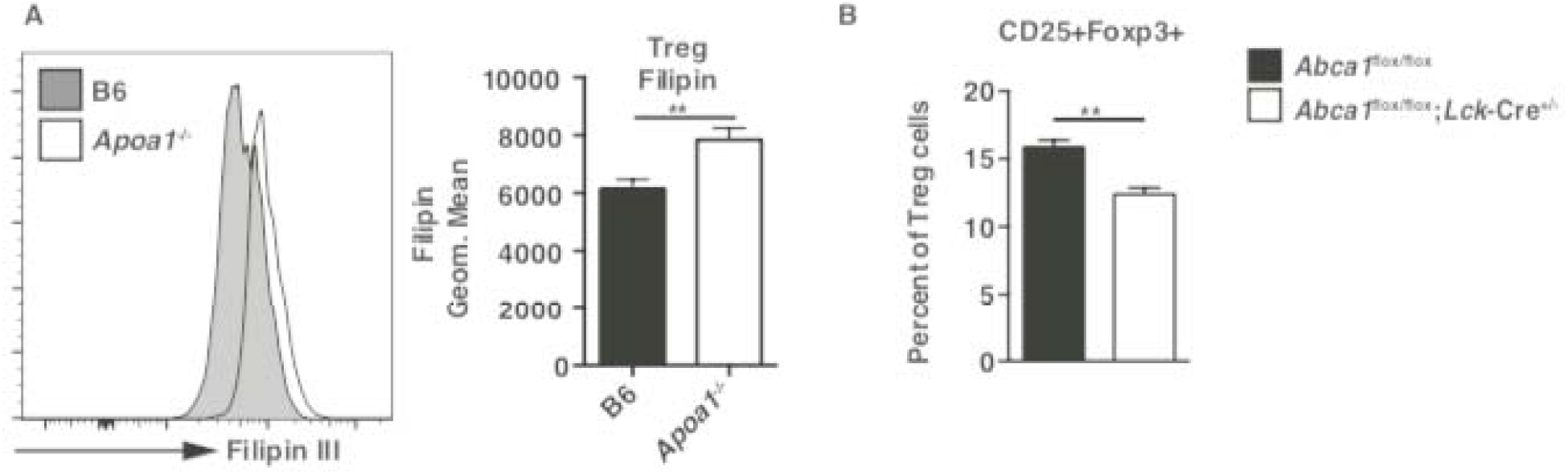
Loss of ApoA1 increases accumulation of cholesterol in Treg cells. **(A)** Intracellular cholesterol content was measured by filipin III staining in B6 and *Apoa1*^−/−^ Treg cells (CD4^+^CD25^+^Foxp3^+^) using flow cytometry. (**B**) PaLNs from *Abca1*^flox/flox^ or *Abca1*^flox/flox^;*Lck*-Cre^+/−^ mice were examined for the percentages of Treg cells (CD4+CD25+Foxp3+) by flow cytometry. Results are shown as representative histogram plots from duplicate experiments and bar graphs represent the mean (± SEM) from two independent experiments (n = 7-10) (**A**) or (n = 4-6) (**B**), \*\**p* < 0.01.

### *Apoa1*^−/−^ Treg cells have a defect in IL-2 signaling

Signaling by IL-2 is essential for the stability and homeostatic proliferation of Treg cells. Therefore, we next checked for defective IL-2 signaling in Treg cells from *Apoa1*^−/−^ mice by examining the levels of phosphorylation of signal transducer and activator of transcription 5 (STAT5), the signaling molecule downstream of the IL-2 receptor. Upon stimulation with IL-2, Treg cells from *Apoa1*^−/−^ showed reduced phosphorylation of STAT5 compared to Treg cells from B6 mice (**Figure 4A**). This defect in IL-2 signaling was not observed in naïve or memory CD4 T cells (**Figure 4B**), suggesting that changes in IL-2 signaling are specifically responsible for the changes observed in Treg cells from *Apoa1*^−/−^ mice.

**Figure 4.**
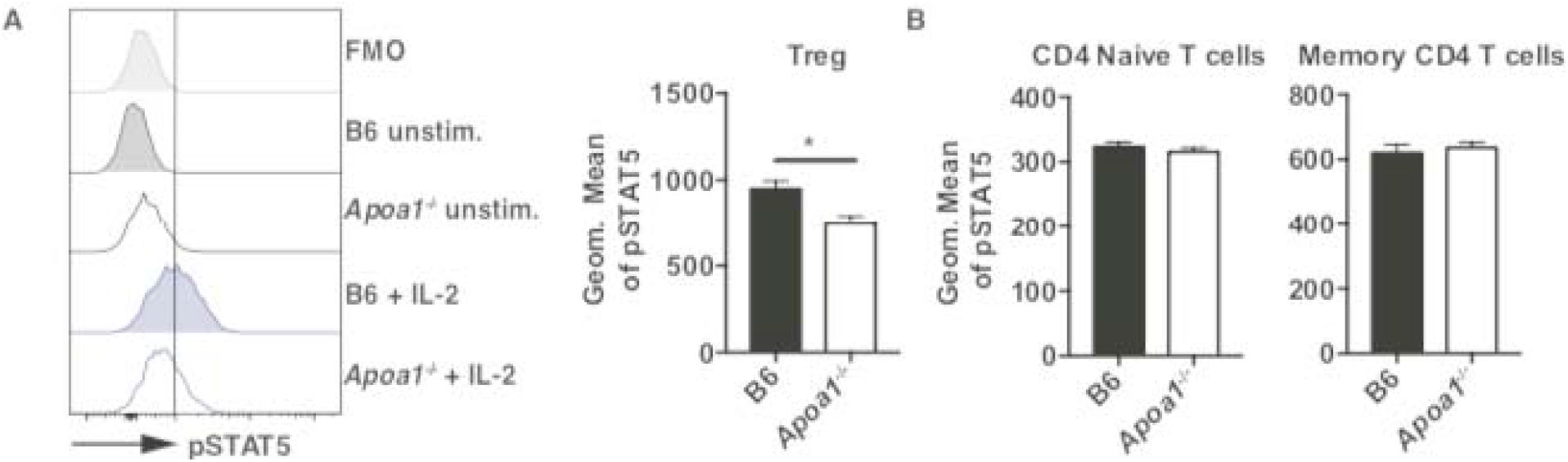
ApoA1 is required for optimal IL-2 signaling in Treg cells. Splenocytes from B6 and *Apoa1*^−/−^ mice were stimulated with IL-2. Treg cells **(A)** as well as naïve and memory CD4^+^ T cells **(B)** were then examined for levels of phosphorylated STAT5. Results are expressed as the mean (± SEM) from one experiment (n = 4-5), \**p* < 0.05.

### IL-2 can rescue *Apoa1*^−/−^ Treg cell induction

We next sought to answer whether induction of naïve T cells to Treg cells was impacted by loss of ApoA1. To accomplish this, naïve CD4 T cells from B6 and *Apoa1*^−/−^ mice were treated with αCD3, αCD28, and TGFβ to derive induced Treg (iTreg) cells *in vitro*. After treatment, fewer iTreg cells were generated from *Apoa1*^−/−^ naïve T cells compared to B6 cells (27.5 ± 4.1 % versus 42.9 ± 2.4 %, respectively) (**Figure 5A**), suggesting that ApoA1 also impacts iTreg generation. When supernatants from these cultures were examined, we found a decrease in the amount of IL-2 produced by cultured Treg cells from *Apoa1*^−/−^ mice compared to those from control mice (**Figure 5B**). This decrease was accompanied by a reduction in both Foxp3 and CD25 (IL-2Rα) expression (**Figure 5C**). Upon adding exogenous IL-2 to culture conditions, we observed a rescue of the percentages of iTreg cells from *Apoa1*^−/−^ naïve T cells to levels comparable with B6 cells (**Figure 5C**). This was also accompanied by increases in both Foxp3 and CD25 levels, supporting the notion that the reduction in Treg cell numbers in *Apoa1*^−/−^ cells is dependent on IL-2 signaling deficits.

**Figure 5.**
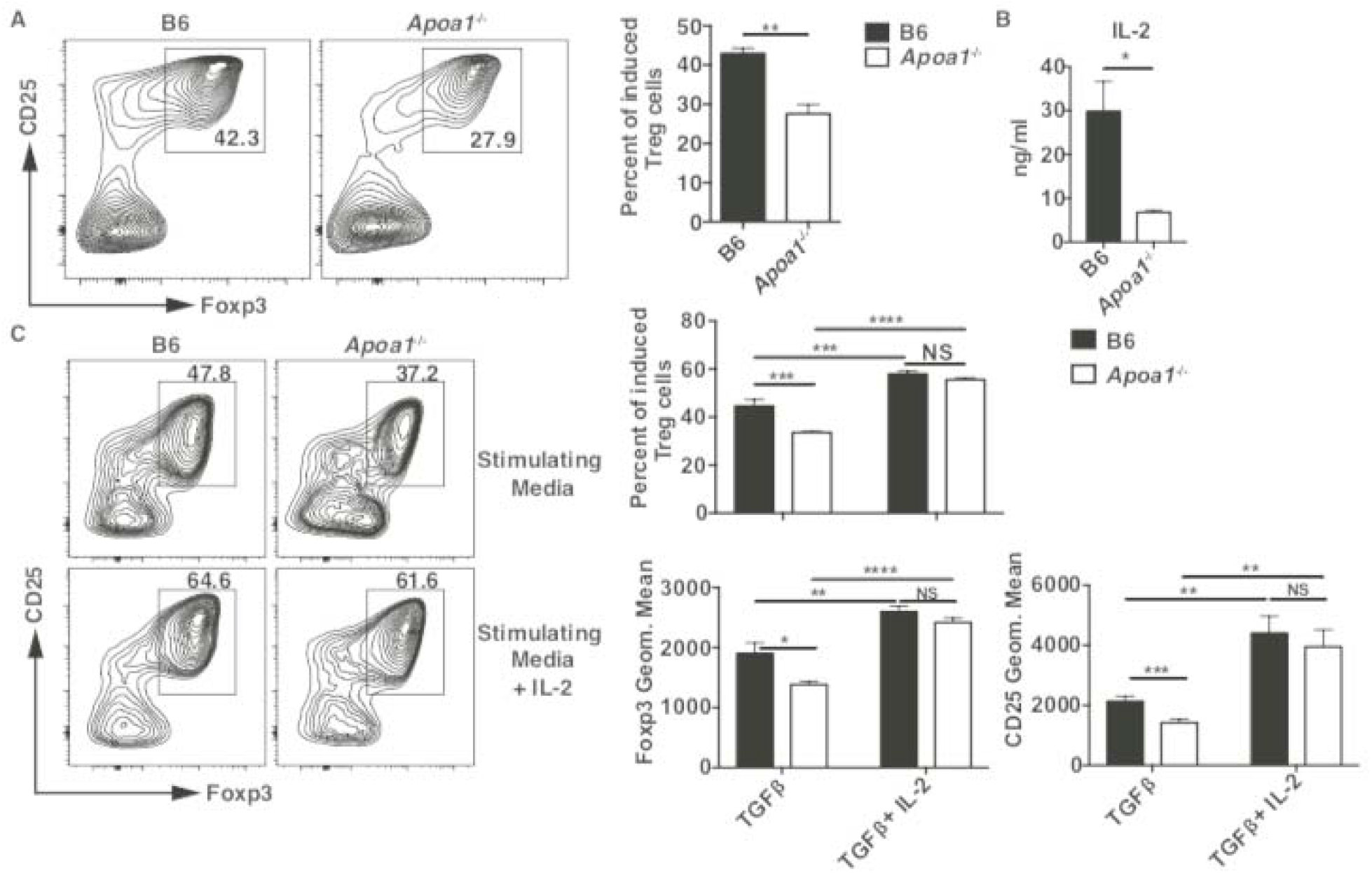
Treg *in vitro* induction is reduced in *Apoa1*^−/−^ naïve T cells but rescued by exogenous IL-2. Naïve (CD25^−^CD44^lo^CD62L^hi^) CD4 T cells sorted from B6 and *Apoa1*^−/−^ mice were stimulated with plate-bound αCD3, soluble αCD28, and TGFβ, without **(A-B)** or with **(C)** exogenous IL-2 for 2 **(B)** or 4 **(A and C)** days. Cells were stained for Foxp3 expression and percentages of induced Treg cells (CD25^+^Foxp3^+^) **(A and C)**, the levels of Foxp3 and CD25 expression on total CD4 T cells **(C)**, and supernatant levels of IL-2 **(B)** were determined by ELISA **(B)** or flow cytometry **(A and C)**. Results are expressed as the mean (± SEM) of triplicate wells from one of two or three independent experiments, \**p* < 0.05, \*\**p* < 0.01, \*\*\**p* < 0.001 and \*\*\*\**p* < 0.0001.

## DISCUSSION

The objective of this study was to determine the role of ApoA1 in Treg cell development during steady state. We found that loss of ApoA1 affects Treg cell homeostasis and function. We observed this defect in both naturally occurring Treg cells as well as induced Treg cells from ApoA1-deficient mice. Importantly, we also found that loss of ApoA1 does not affect either naïve effector or memory CD4 T cells. A number of laboratories including our own have shown that administration of ApoA1 protects against Treg cell loss and conversion of Treg cells to inflammatory T cells during atherosclerosis progression (7–9). In addition, ApoA1 has been shown to reduce inflammation as well as lymphocyte involvement in inflammatory conditions such as rheumatoid arthritis and lupus (11, 12). In this report, our results show that ApoA1 is inherently beneficial to Treg cell homeostasis, regardless of the atherosclerotic or inflammation status of the host.

We found an accumulation of cholesterol in Treg cells from ApoA1-deficient mice, suggesting that lack of ApoA1 causes disruptions in normal Treg cell cholesterol homeostasis. This is observation is consistent with the previous finding that chimeras with loss of ABCA1 exhibit a similar defect in Treg cell homeostasis. Furthermore, these results are in agreement with work from our laboratory which demonstrates that loss of the cholesterol transporter ATP-binding cassette subfamily G member 1 (ABCG1) impacts both T cell and thymic and peripheral Treg cell development (16) (17).

The results presented here show that lack of ApoA1 causes perturbations in IL-2 receptor signaling and STAT-5 phosphorylation. Lipid rafts in the plasma membrane play an important role in T cell functioning (18). Since IL-2 receptor signaling occurs at the plasma membrane on lipid rafts, changes in plasma membrane cholesterol composition could affect the signaling ability of the IL-2 receptor. In fact, minor changes in cholesterol content and plasma membrane lipids can alter IL-2 signaling events (19, 20) and subsequently affect phosphorylation of STAT5. IL-2 is essential for Treg cell expression of Foxp3, a transcription factor critically involved in Treg cell development (21). Concordant with this, here we showed that addition of exogenous IL-2 increased levels of Foxp3 protein expression and rescued defects in Treg cell induction from *Apoa1^−/−^*, naïve T cells *in vitro*. We have previously reported that administration of ApoA1 in atherosclerotic mice can increase IL-2Rα levels on Treg cells (9). Thus, our results indicate that ApoA1 regulation of IL-2/IL-2R signaling can occur at steady state and in the absence of atherosclerosis.

The benefits of ApoA1 as a protective, anti-inflammatory agent and as an immune system regulator have been well documented (10, 11, 13, 22, 23). Lack of ApoA1 increases T cell proliferation and release of inflammatory cytokines IFN-γ and IL-17 in a rheumatoid arthritis animal model (12). Similarly, *Apoa1* transgenic mice, which display increased levels of ApoA1, are protected against lupus and show reduced lymphocyte activation (11). A more recent study examining the role of ApoA1 binding protein (AIBP) showed that inhalation of AIBP lowered lipopolysaccharide (LPS)-induced airway neutrophilia and lung injury in mice (24). These effects are seen outside of the protective role of ApoA1 in the cardiovascular system, suggesting that the benefits of ApoA1 extend beyond heart disease. Indeed, our results demonstrate that ApoA1 can act as an immune modulator in inflammatory disease, helping increase the protective Treg cell population and putting a check on activated effector T cells. Whether infusion of lipid-free ApoA1 can affect the percentages of Treg cells in atherosclerosis is yet to be determined. Finally, whether or not increased levels of ApoA1 are protective against other inflammatory conditions is unknown. Future work focused on answering these questions may lead to improved therapeutics for inflammatory conditions such as atherosclerosis.

## ACKNOWLEDGEMENTS

We thank Debbi Yoakum for assistance with mouse colony management and the La Jolla Institute (LJI) Flow Cytometry Core Facility for cell sorting. This project was supported by the American Heart Association (13POST16990031 to D.E.G.), the American Diabetes Association (#7-12-MN-13 to D.E.G. and C.C.H.), and the National Institutes of Health (NHLBI P01 HL055798 to C.C.H, R01 HL112276 to C.C.H. and M.S.T., R01 HL127649 to M.S.T, and S10 RR027366-01A1 to LJI Flow Cytometry Core Facility). This content is solely the responsibility of the authors and does not necessarily represent the official views of the National Institutes of Health.

